## Supplementary Figures 1 - 5 for "MolNetEnhancer: enhanced molecular networks by integrating metabolome mining and annotation tools"

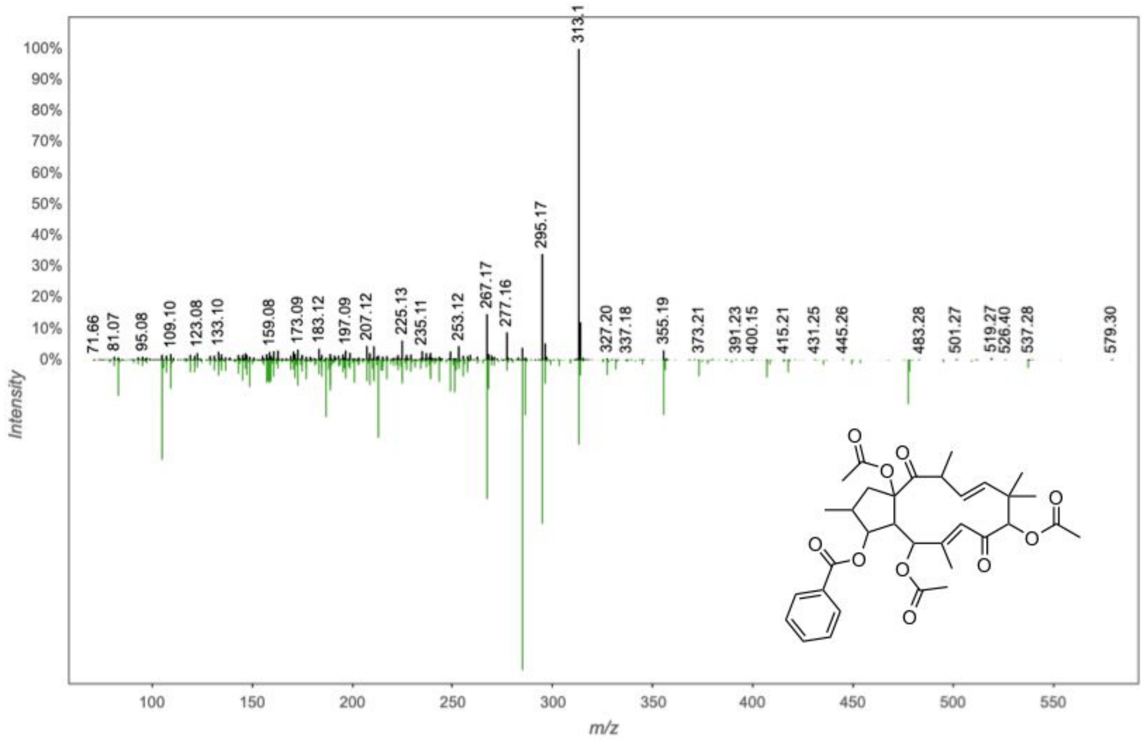


**Supporting Figure S1**. Mirror plot comparing molecular feature with *m/z* 614.30 and RT 373.17 (black) to GNPS reference spectrum of a jatrophane diterpenoid (green). A total of 289 shared peaks were found. Mass peaks at *m/z* 313, 295, 285 are characteristic for a *Euphorbia* diterpenoid backbone skeleton, however spectral similarity (cosine score) was only found to be 0.71. The unknown molecular feature is thus likely a close structural analogue of the jatrophane diterpenoid. The GNPS reference spectrum as well as the mirror plot is publicly accessible at https://gnps.ucsd.edu/ProteoSAFe/result.jsp?task=26326c233918419f8dc80e8af984cdae&view=view_all_annotations_DB#%7B%22main.%23Scan%23_lowerinput%22%3A%223633%22%2C%22main.%23Scan%23_upperinput%22%3A%223633%22%7D.

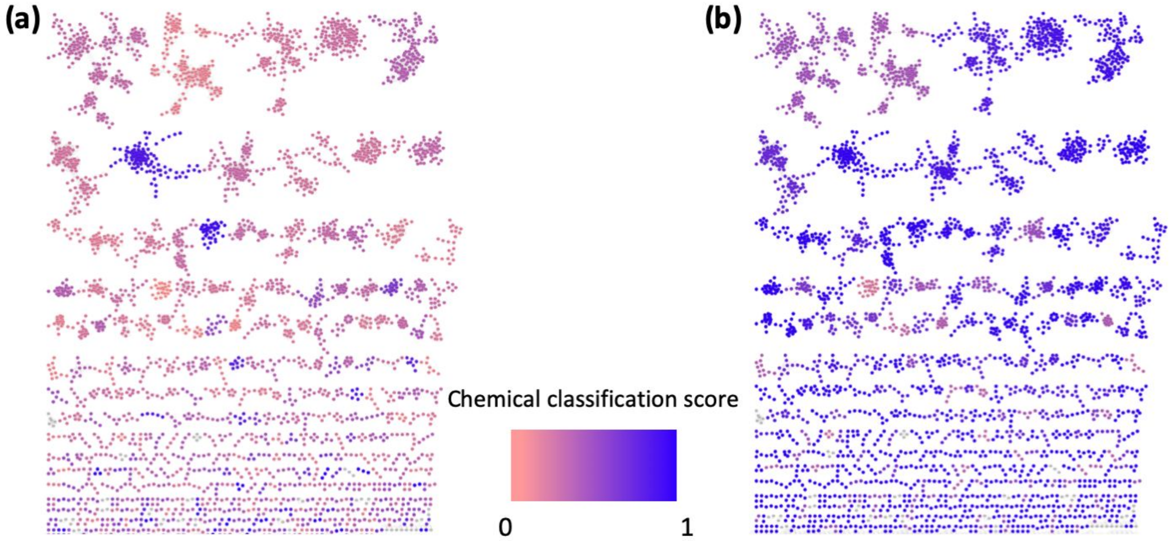


**Supporting Figure S2**. (**a**) Marine sediment Salinispora/Streptomyces molecular network colored by chemical classification scores for annotated chemical class terms, and (**b**) same molecular network colored by chemical classification scores for annotated chemical kingdom terms. Light grey means no database matches were found. The higher the class score, the more consistent the chemical annotations are. The kingdom scores represent the database coverage of nodes across a molecular family with scores closer to zero representing families with fewer nodes that have at least one database hit. Whilst most MFs do have database matches for all or most nodes, the consistency in chemical class annotations is - apart from some exceptions - less (indicated by the more orange/pink colors in the left panel). This indicates that for many MF family members the right molecular structures might not yet be present in the structural databases used.

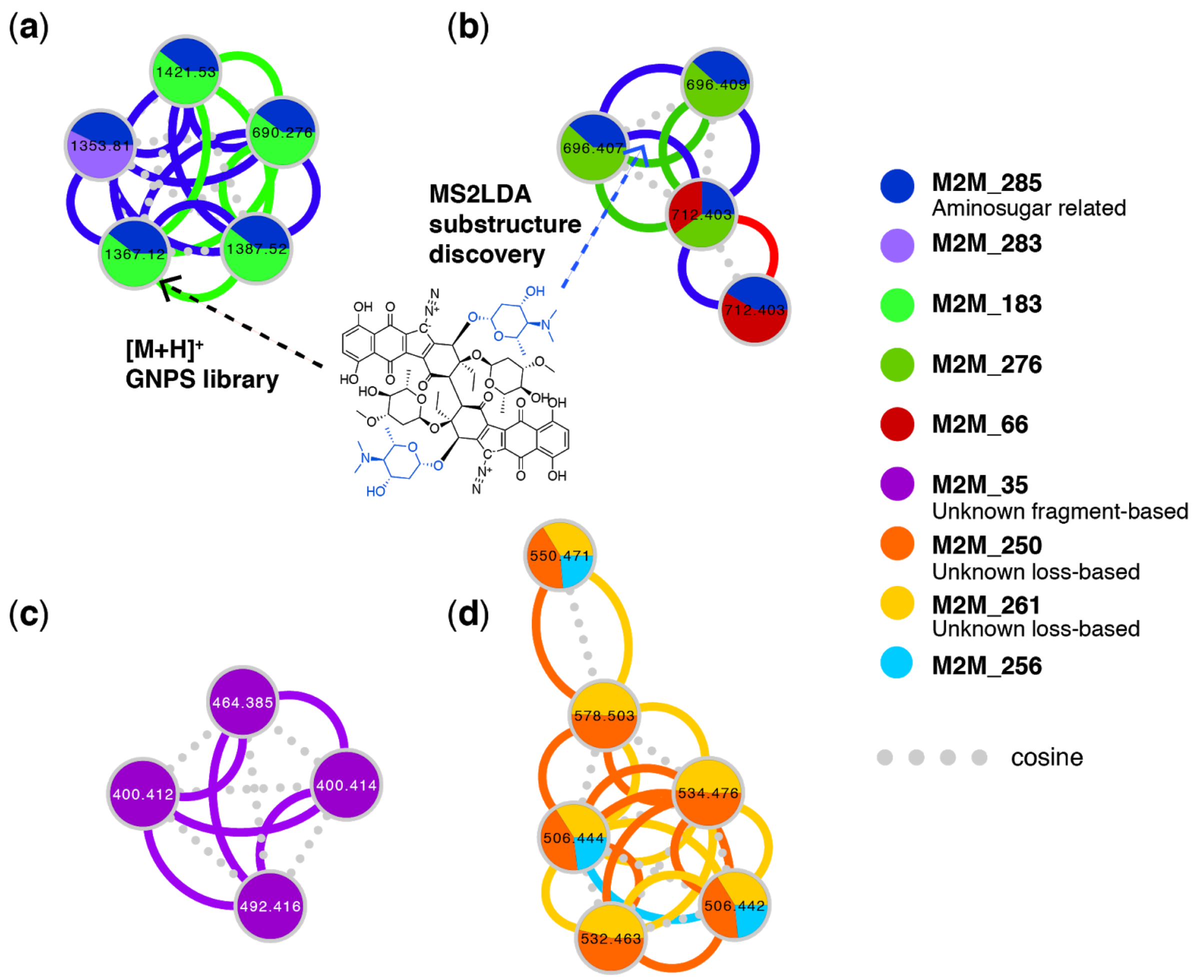


**Supporting Figure S3**. Molecular families from marine sediment bacteria enriched with chemical information by color coded Mass2Motif substructure information mapped on them, with (**a**) lomaiviticin related molecular family found through GNPS library hits where all members share the amino sugar related motif as coloured blue in the depicted lomaiviticin A structure, (**b**) yet unknown molecular family that shares the same amino sugar related motif as highlighted in (a), (**c**) yet unknown molecular family sharing an unknown fragment-based motif occurring 0.7% in the marine sediment data set, and (**d**) yet unknown molecular family sharing unknown loss-based motifs occurring 0.4% (Mass2Motif 250) and 0.8% (Mass2Motif 261) in the marine sediment data. In all MFs, nodes are coloured based on motif overlap scores and the edges present similar colours to show if cosine score-connected nodes share similar Mass2Motifs. It can be seen that in most families multiple motifs are shared across some of its members.

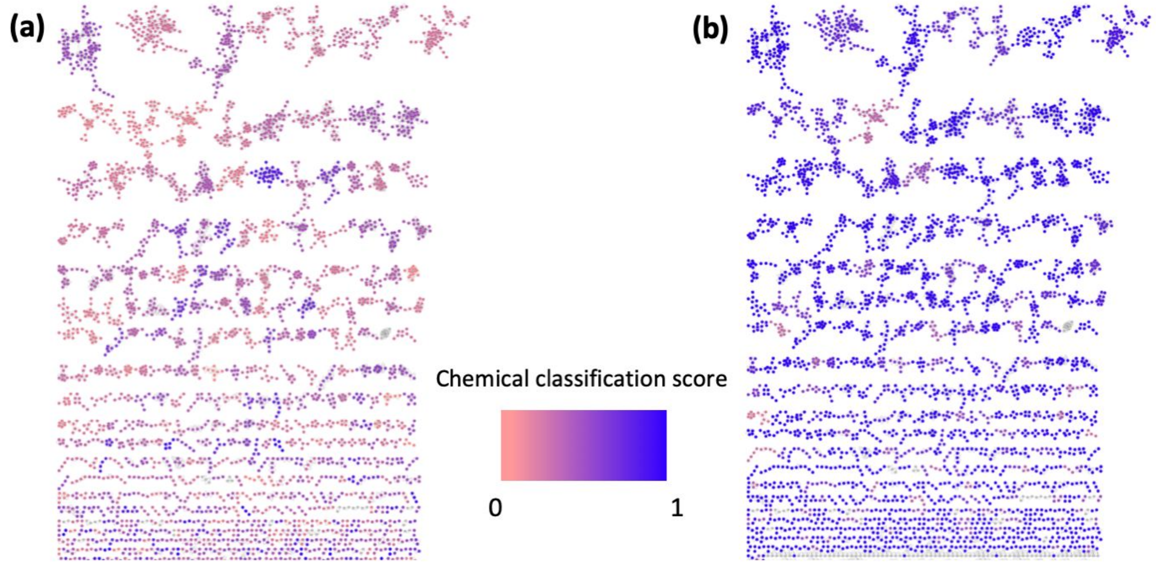


**Supporting Figure S4**. (**a**) Nematode symbionts *Photorhabdus/Xenorhabdus* network colored by chemical classification scores for annotated chemical class terms, and (**b**) same molecular network colored by chemical classification scores for annotated chemical kingdom terms. Light grey means no database matches were found. The higher the class score, the more consistent the chemical annotations are. The kingdom scores represent the database coverage of nodes across a molecular family with scores closer to zero representing families with fewer nodes that have at least a database hit. We observe database coverages of close to 1 for most molecular families; however, some molecular families have a lower coverage with a few nodes that return candidate structures. Furthermore, we observe that the chemical class annotation is not always consistent indicating that manual inspection and validation of those hits remains essential.

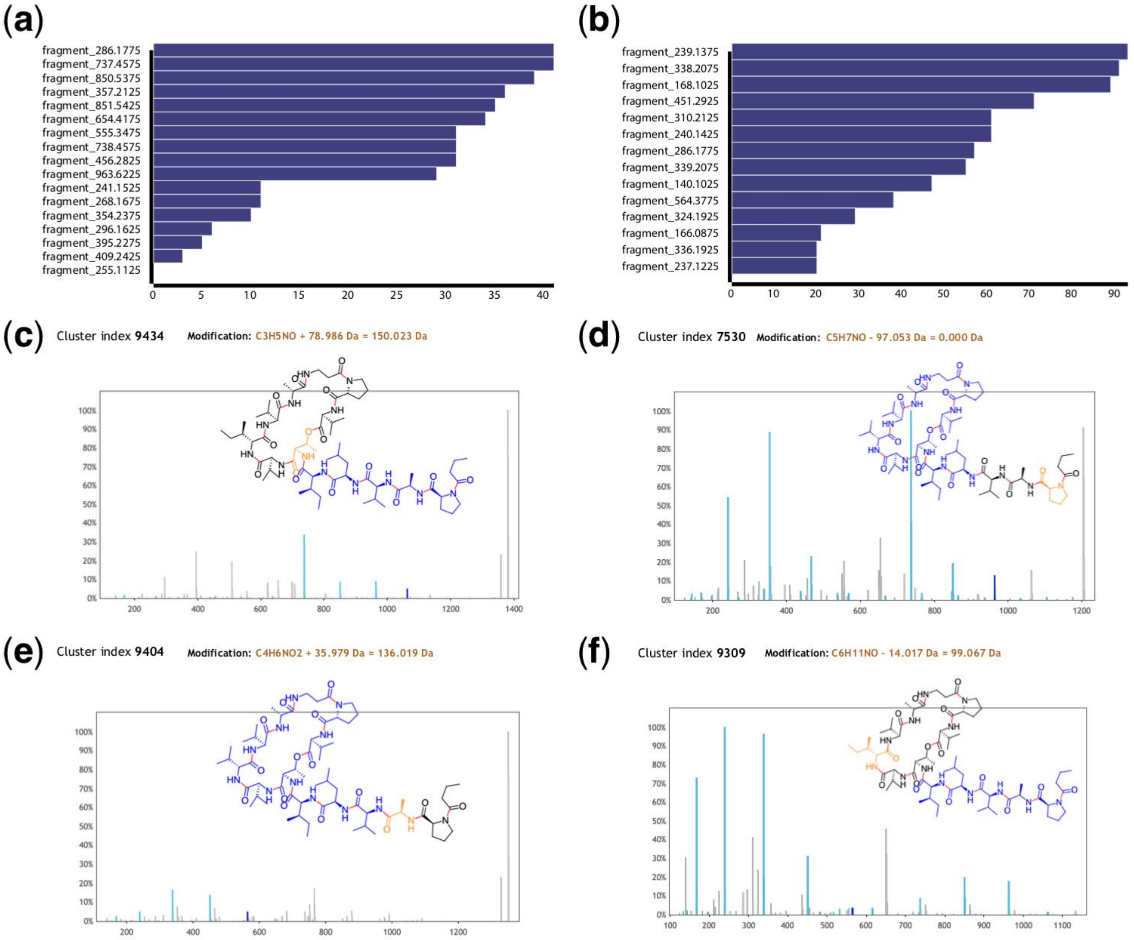


**Supporting Figure S5**. Xenoamicin Mass2Motif mass feature frequency plots for (a) Mass2Motif related to xenoamicin peptidic ring and (b) xenoamicin peptidic tail. It can be observed that many mass fragments are present in at least 75% of the associated molecular features (9 and 6 for ring and tail Mass2Motif, respectively) with a few mass fragments present in nearly all associated molecular features. (c) and (d) Examples of annotated xenoamicin A modified structures in which only the ring Mass2Motif was found. Indeed, we observe that VarQuest annotates a modified amino acid (addition and loss of) in the tail region of xenoamicin A indicated in orange. (e) and (f) Examples of annotated xenoamicin B modified structures in which only the ring Mass2Motif was found. Indeed, we observe that VarQuest annotates a modified amino acid (double water addition, loss of methyl) in the ring region of xenoamicin B indicated in orange. The structures of xenoamicin A and B differ in one methyl group on the amino acid highlighted in orange in (f) where B has an isobutyl group and A an isopropyl group. In fact, the structure of xenoamicin A is correctly annotated by VarQuest to this fragmented doubly charged ion.
